# ACKR3 Proximity Labeling Identifies Novel G protein- and β-arrestin-independent GPCR Interacting Proteins

**DOI:** 10.1101/2024.01.27.577545

**Authors:** Chloe Hicks, Julia Gardner, Dylan Scott Eiger, Nicholas D. Camarda, Uyen Pham, Saisha Dhar, Hailey Rodriguez, Anand Chundi, Sudarshan Rajagopal

## Abstract

The canonical paradigm of GPCR signaling recognizes G proteins and β-arrestins as the two primary transducers that promote GPCR signaling. Recent evidence suggests the atypical chemokine receptor 3 (ACKR3) does not couple to G proteins, and β-arrestins are dispensable for some of its functions. Here, we employed proximity labeling to identify proteins that interact with ACKR3 in cells devoid of β-arrestin. We identified proteins involved in the endocytic machinery and evaluated a subset of proteins conserved across several GPCR-based proximity labeling experiments. We discovered that the bone morphogenic protein 2-inducible kinase (BMP2K) interacts with many different GPCRs with varying dependency on β-arrestin. Together, our work highlights the existence of modulators that can act independently of G proteins and β-arrestins to regulate GPCR signaling and provides important evidence for other targets that may regulate GPCR signaling.

**Graphical Abstract:** 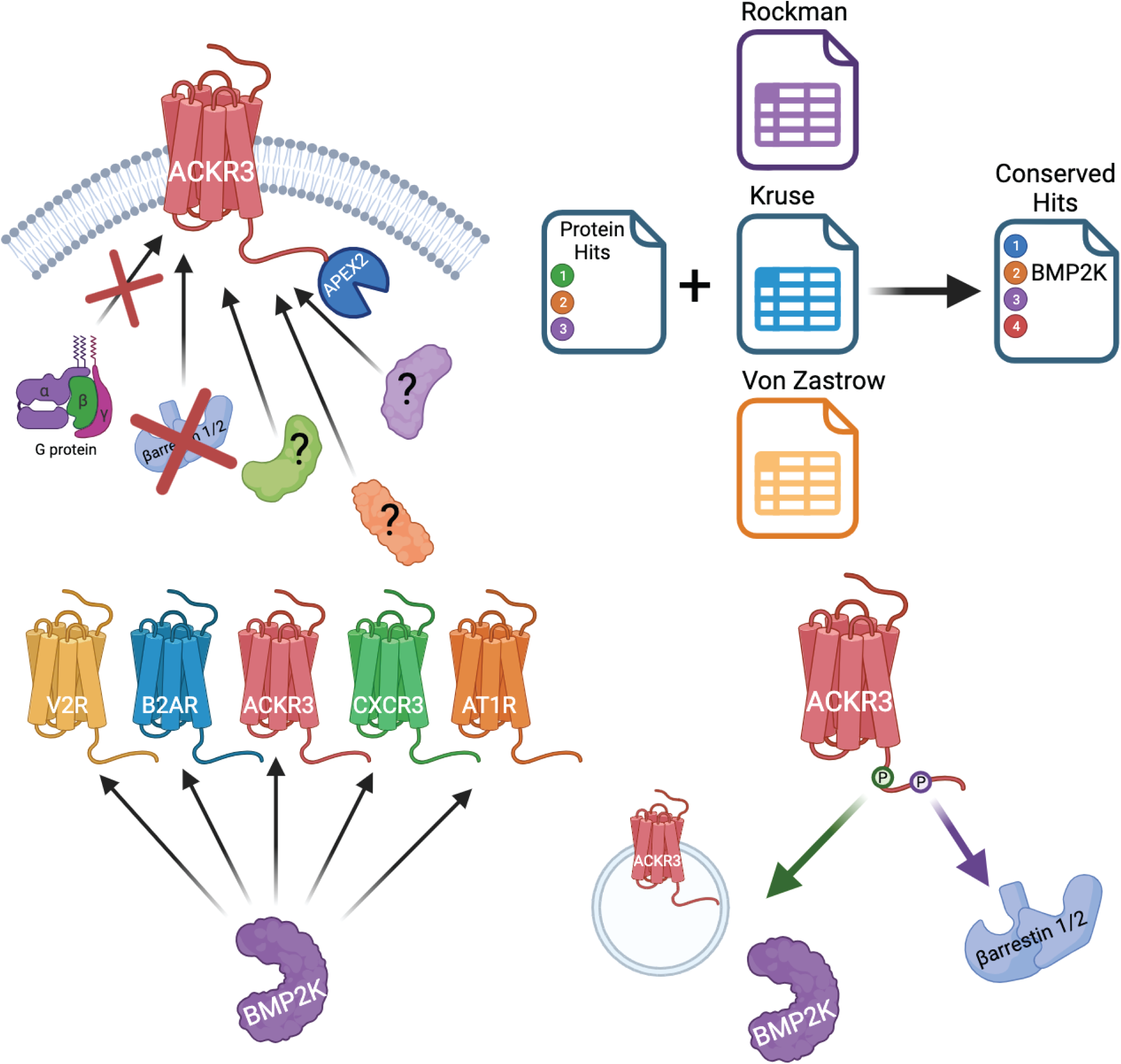

## INTRODUCTION

G protein-coupled receptors (GPCRs) are the largest family of receptors in the human body, and represent the target of approximately one-third of all FDA-approved pharmaceutical drugs (*1, 2*). Most GPCRs signal through two main transducers: heterotrimeric G proteins and β-arrestins. In the canonical GPCR signaling paradigm, a ligand binds to the receptor’s extracellular domain, causing a conformational change that allows for heterotrimeric G proteins to engage with the receptor (*3–5*). G protein activation involves the catalytic exchange of GDP for GTP, resulting in dissociation of the Gα subunit from the Gβγ subunits. Gα subunits modulate effector proteins that regulate levels of downstream signaling molecules like cyclic AMP (cAMP), diacylglycerol (DAG) and inositol 1,4,5-trisphosphate (IP3) (*6, 7*). Following G protein activation, the GPCR Kinases (GRKs) phosphorylate the intracellular surface of the receptor, promoting the recruitment of β-arrestins (*8*). β-arrestins desensitize G protein signaling by sterically hindering G protein coupling, and promote receptor internalization by scaffolding endocytic proteins, such as clathrin and the adaptor protein 2 (AP2) (*9, 10*). β-arrestins can also regulate receptor signaling processes independently of G proteins by activating key signaling molecules, including mitogen-activated protein kinases (MAPKs), non-receptor tyrosine kinase Src, and the protein kinase B (PKB/Akt) (*11–13*).

Given the ubiquity of GPCRs in most cell types and their implication in many pathologies, considerable effort has focused on elucidating the various mechanisms underlying GPCR activation and subsequent signaling responses. As a result, G proteins and β-arrestins are now recognized as canonical mediators of GPCR signaling. Interestingly, certain ligands or cellular conditions can induce preferential activation of G protein or β-arrestin pathways downstream of a receptor, a phenomenon dubbed *biased signaling* (*14, 15*). However, given the specificity and complex cellular responses initiated by GPCRs, sophisticated mechanisms that fine-tune downstream signaling beyond activating two signaling transducers are likely required. Researchers have found that GPCRs can interact with a variety of other GPCR interacting proteins (GIPs), like the receptor activity-modifying proteins (RAMPs) and Homer proteins, which can regulate receptor trafficking and activity (*16–19*). However, it is unclear if the activity of GIPs is always dependent on G proteins and/or β-arrestins, or if there are G protein- and β-arrestin-independent effectors. *Noncanonical GPCR signaling* includes both the novel roles of G proteins and β-arrestins beyond their classically described functionality, and other modulators of GPCR signaling outside of these two canonical transducers (*20–27*). Identifying GPCR signaling partners that regulate signaling independently of G protein or β-arrestin could provide novel therapeutic drug targets.

Studying G protein- and β-arrestin-independent GPCR signaling pathways is difficult given the near ubiquitous requirement of these proteins for proper receptor function. However, the atypical chemokine receptor 3 (ACKR3), formerly known as the C-X-C chemokine receptor type 7 (CXCR7), provides a useful opportunity to study noncanonical GPCR signaling. ACKR3 was previously classified as a scavenger receptor for chemokines CXCL11 and CXCL12, as its binding of these ligands did not activate G proteins (*28–31*). However, it was later discovered that ligand binding to ACKR3 recruits β-arrestin, leading to the reclassification of ACKR3 as a completely β-arrestin-biased receptor (*32, 33*). It is hypothesized that the lack of G protein activation at ACKR3 is due to alterations in the canonical DRY motif of the receptor’s second intracellular loop (ICL2), which is crucial for G protein coupling (*34, 35*). Interestingly, recent evidence demonstrates that β-arrestins are dispensable for some functionality at ACKR3. For example, receptor phosphorylation, but not β-arrestin, is necessary for ACKR3 to internalize, promote interneuron motility, and regulate CXCL11 and CXCL12 levels (*36, 37*). These findings suggest that noncanonical signaling mechanisms that promote GPCR signaling independently of both G proteins and β-arrestins may exist.

ACKR3, as a receptor that (1) does not activate G proteins, (2) recruits β-arrestin, but (3) demonstrates functionality independent of β-arrestin, serves as a unique model system to identify noncanonical GPCR signaling partners. Prior studies have successfully demonstrated the use of proximity labeling to interrogate protein:GPCR interactions at different receptors, using different ligands, and even at different subcellular locations (*38–43*). These prior studies have laid the foundation for our understanding of the GPCR “interactome”. Here, we performed proximity labeling of the β-arrestin-biased receptor ACKR3 in cells devoid of β-arrestin to determine what proteins may interact with a GPCR independently of G proteins and β-arrestins (*39–41*). We identified ∼100 proteins whose interactions with ACKR3 significantly changed following receptor activation, of which many were involved in mechanisms of endosomal trafficking and recycling. We compared our data to three other published GPCR-based proximity labeling datasets, where G protein and β-arrestin signaling were preserved, and identified 17 conserved proteins. Gene Ontology and subsequent pathway analyses revealed that these effectors are likely involved in receptor endocytosis. Finally, we explored the role of one conserved interacting partner, bone morphogenic protein 2-inducible kinase (BMP2K), in GPCR signaling. Using various biochemical assays, we determined that BMP2K recruits to ACKR3 in a β-arrestin-independent fashion, but also recruits to a variety of other therapeutically-relevant GPCRs, suggesting that it may be a highly conserved GIP. We provide concrete evidence for G protein- and β-arrestin-independent GPCR interacting partners that may serve as novel druggable targets for modulating GPCR signaling activity.

## RESULTS

### APEX2 Proximity Labeling of ACKR3 in β-arrestin-1/2 knockout HEK293 cells reveals noncanonical GPCR signaling effectors

While many GPCRs require β-arrestin for receptor internalization, recent studies have demonstrated ACKR3 can internalize in the absence of β-arrestin, but in a manner requiring receptor phosphorylation (*36, 37*). We evaluated ACKR3 trafficking to early endosomes in wild-type (WT) HEK293, β-arrestin-1/2 knockout (KO), and GRK-2/3/5/6 KO cell lines using Bioluminescence Resonance Energy Transfer (BRET). Consistent with previous data, ACKR3 internalized independent of β-arrestins, regardless of the endogenous (CXCL12) or synthetic (WW36 and WW38) ligand used to stimulate the receptor (**Figure 1A-1D**). Interestingly, we detected ACKR3 internalization in the absence of GRKs, with statistically insignificant increases in internalization levels upon rescue of each GRK isoform (**Figure 1E**). These results build upon previous studies showing that β-arrestins are dispensable in ACKR3 internalization by also demonstrating that GRKs are not required to drive ACKR3 internalization (*44, 45*).

**Figure 1:**
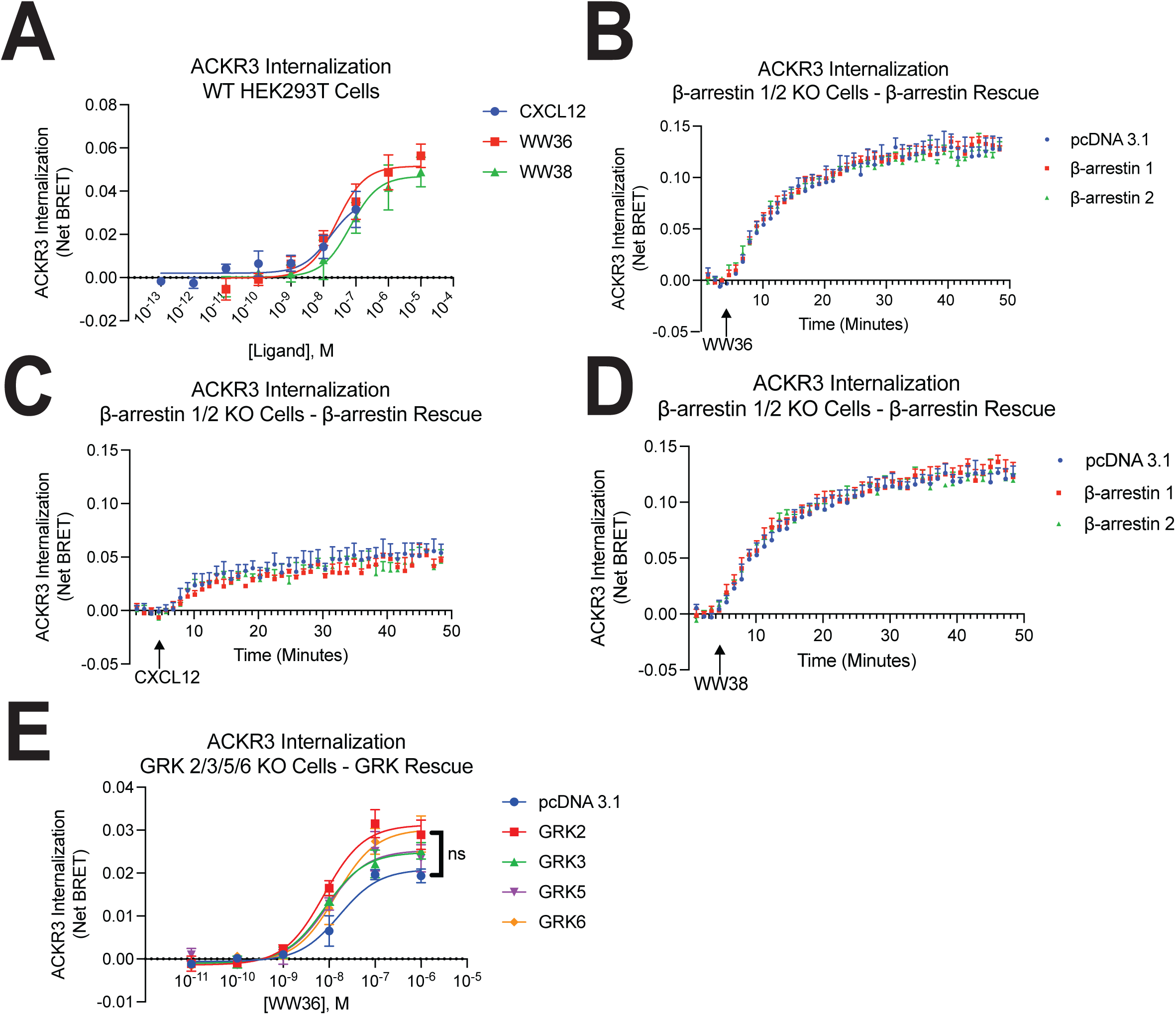
ACKR3 endocytosis does not require *β*-arrestins or GRKs. **(A)** Agonist concentration-response at thirty minutes of ACKR3 trafficking to early endosomes in HEK293 cells. Data shown are the mean ± SEM of n=5. Kinetic time-course of ACKR3 trafficking to early endosomes following stimulation with (**B**) WW36, (**C**) CXCL12, and (**D**) WW38 in β-arrestin 1/2 knockout cells with β-arrestin rescue. Data shown are the mean ± SEM of n=3. (**E**) Agonist concentration-response at thirty minutes of ACKR3 trafficking to early endosomes in GRK 2/3/5/6 knockout cells with and without addback of each GRK isoform. Data shown are the mean ± SEM of n=4. Significance testing was performed using one-way ANOVA comparing the net BRET ratio at maximum dose between empty vector and other transfection conditions (pcDNA 3.1 vs. GRK).

To identify G protein- and β-arrestin-independent GPCR effectors, we generated an ACKR3 receptor construct with a C-terminal modified ascorbate peroxidase enzyme (APEX2). Following incubation with biotin phenol, cells were treated with hydrogen peroxide (H_2_O_2_) to catalyze a one-electron transfer reaction between APEX2 and biotin phenol, generating biotin-phenoxyl radicals that covalently bind to endogenous proteins within a 20-nm proximity to the APEX2 enzyme **(Figure 2A)** (*46, 47*). Biotinylated proteins are then isolated and identified using mass spectrometry.

**Figure 2:**
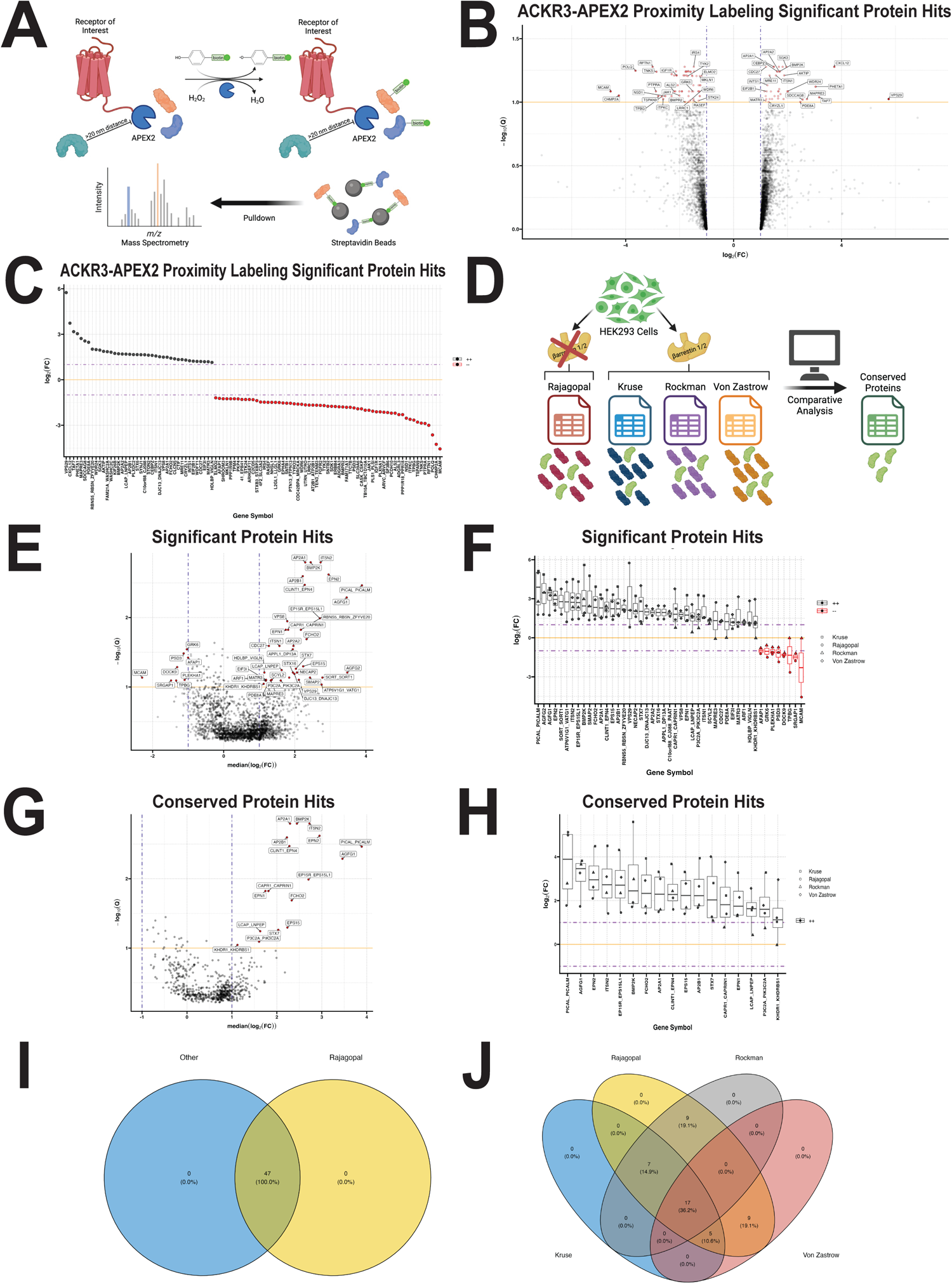
ACKR3 proximity labeling and comparison of APEX datasets reveal conserved GPCR interacting proteins. **(A)** Schematic of ACKR3-APEX2 proximity labeling technique. **(B)** Volcano and **(C)** waterfall plots showing proteins with significant fold changes in proximity to ACKR3 following stimulation with CXCL12. Significance was determined by calculating the log 2-fold change and identifying those with a fold change greater than 2 or less than 0.5 and a False Discovery Rate (FDR)-corrected combined Fisher p-value cutoff of 0.1. ACKR3-APEX2 proximity labeling protein hits are representative of n=3. (**D**) Schematic design of comparative analysis used to identify conserved GPCR interacting proteins. **(E)** Volcano and **(F)** waterfall plots showing all proteins identified in the four datasets with significant fold changes in interaction with a GPCR following activation. Proteins identified in at least 2 of the 4 datasets are shown (significant protein hits). **(G)** Volcano and **(H)** waterfall plots showing the proteins identified in every dataset (conserved protein hits). Significance was determined by calculating the median fold change for each protein across every dataset that included the protein. Identified proteins exhibited a median fold change greater than 2 or less than 0.5 and had a False Discovery Rate (FDR)-corrected combined Fisher p-value cutoff of 0.1. **(I)** Venn diagram showing overlap in the significant protein hits found in the present dataset (Rajagopal) and at least one of the other three GPCR APEX datasets. **(J)** Multi-set Venn diagram showing the number of significant protein hits identified in each combination of the four GPCR APEX datasets.

We generated and validated β-arrestin 1/2 KO HEK293 cells stably expressing ACKR3-APEX2 (**Supplemental Figures 1A-1F)** (*48*). HEK293 cells were used due to their lack of endogenous ACKR3 expression and prior use in other GPCR proximity labeling assays (*49*). Our experimental design allowed us to probe potential protein interactions of a β-arrestin-biased receptor in a cellular system devoid of β-arrestin. We stimulated these cells with ACKR3’s endogenous ligand CXCL12 or vehicle control, and then isolated biotinylated proteins for subsequent analysis with mass spectrometry to identify potential ACKR3-interacting proteins (**Supplemental Figure 1G**).

Of the nearly 6000 proteins identified using mass spectrometry, 101 showed a significant fold change in proximity with ACKR3 following stimulation with the endogenous ligand CXCL12, 40 of which showed an increase and 61 of which showed a decrease (**Figure 2B-2C**). Many of the proteins with increased ACKR3 proximity are involved in mechanisms of endosome formation, trafficking, sorting, and recycling. For instance, the protein with the greatest positive fold change is the vacuolar protein sorting-associated protein 29 (VPS29), a component of the retromer and retriever complexes, which is involved in sorting protein cargo from endosomes to the trans-Golgi network and the plasma membrane (*50–52*). Protein components implicated in other cargo sorting complexes, such as the adaptor protein 2 (AP2) and FTS-Hook-FHIP (FHF) complex were identified as well, suggesting G protein- and β-arrestin-independent mechanisms for activating this endocytic machinery (*50, 53, 54*). Other identified proteins are regulators of microtubule binding activity, suggesting that ACKR3 may undergo endosomal sorting and trafficking using microtubules (*55*). Many of the 61 protein hits with decreased ACKR3 interactions are plasma membrane-localized proteins, potentially suggesting that, following CXCL12 stimulation, ACKR3 undergoes endocytosis and is trafficked to subcellular compartments.

### Comparison of different GPCR interactomes reveals conserved canonical and noncanonical GPCR effectors

An inherent limitation in proximity labeling is the identification of biologically inconsequential proteins due to non-specific binding of proteins during purification, bystander effects, and endogenous biotinylation of proteins, among others (*56–58*). To more confidently identify biologically meaningful GIPs, we selected three other GPCR-based proximity labeling datasets performed in HEK293 cells for further analyses (**Figure 2D)**. These studies, conducted in the laboratories of Andrew Kruse, Howard Rockman, and Mark von Zastrow, identified potential protein-protein interactions at various GPCRs, including the β_2_-adrenergic receptor (β_2_AR), angiotensin II type I receptor (AT_1_R) and the δ-opioid receptor (DOR) (*38–40*). We performed two major comparative analyses: (1) we identified proteins whose abundance (i.e., proximity) changed significantly in the same direction across at least two of the four data sets **(Figure 2E-2F)** and (2) those that changed significantly in the same direction across all four data sets **(Figure 2G-2H).** Together, these approaches strengthen our ability to identify meaningful GIPs, at the cost of potentially removing less statistically powerful but biologically significant potential interactions.

We identified 47 proteins that showed significant changes in their interactions with a GPCR following receptor activation in at least two out of four data sets (**Figure 2E-2F**). Of these interactions, 39 proteins experienced increased fold changes while 8 had decreased changes. We also performed an analysis excluding the Kruse dataset, as it contained a single replicate, but found its exclusion had little impact on the proteins identified (**Supplemental Figure 2A-2D**). Of the 47 proteins identified in at least two of the datasets, 17 were conserved across all four datasets, and all 47 proteins were present in our ACKR3 based experiment (**Figure 2G-2I)**. More specifically, we found that, aside from the 17 proteins identified across all four datasets, 12 were identified in three datasets, and 18 were found in ours and one other dataset (**Figure 2J**). Together, these analyses reflect a set of highly conserved potential protein:GPCR interacting partners that may vary in their dependence on G protein or β-arrestin functionality.

### Pathway analyses reveal clathrin-dependent endocytosis as a prominent conserved biological process that variably depends on G proteins and β-arrestins

We performed Gene Ontology analyses of the 17 conserved protein hits to assess their reported molecular function, cellular compartment, and biological processes (**Figure 3A-3C**) (*59*). We found that many of these proteins localize to the plasma membrane or clathrin-coated pits and vesicles and have roles in clathrin binding, membrane binding, and endocytic adaptor activity (**Figure 3A-3B**). The top biological processes identified involve endocytosis and/or endosomal transport (**Figure 3C**). Further, several of the conserved proteins are also implicated in intracellular trafficking, including transport from the Golgi to endosomes and vesicle recycling. Endocytosis was the most prominent pathway in the Kyoto Encyclopedia of Genes and Genomes (KEGG) enrichment analysis. Various infectious processes were also identified, which may suggest the involvement of these particular endocytic mechanisms in bacterial and viral entry into the cell (**Figure 3D**) (*60*). Other implicated cellular pathways involve maintaining cellular structural integrity, such as through cell-cell junctions and cytoskeleton regulatory proteins. Ultimately, our data demonstrates that receptor-mediated endocytosis, particularly clathrin-mediated, is a highly conserved mechanism in GPCR signaling. The implicated proteins are likely conserved at many GPCRs, but their dependence on G proteins and/or β-arrestins is not clear.

**Figure 3:**
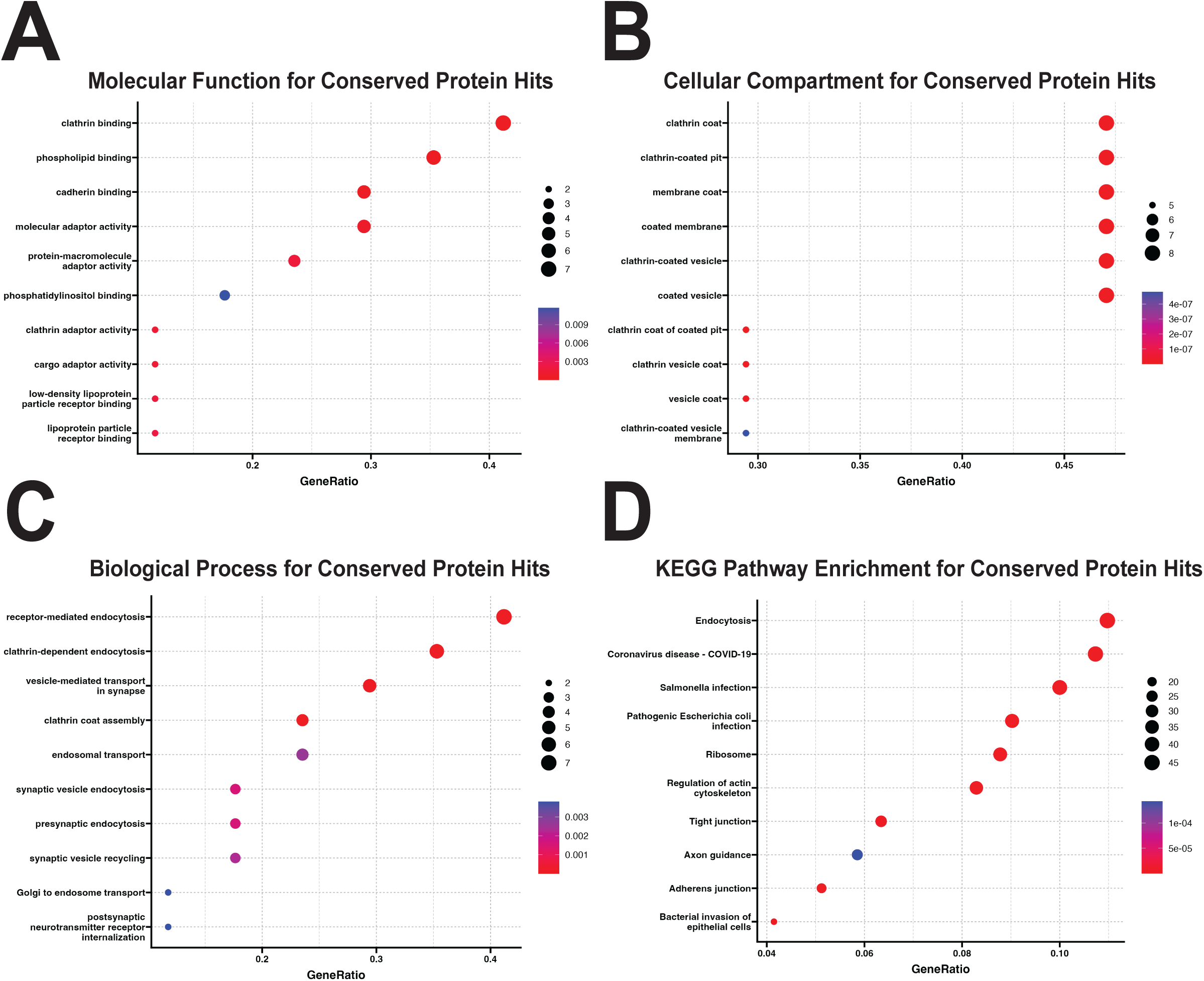
Gene Ontology and KEGG pathway enrichment for conserved GPCR interacting proteins identify clathrin binding and receptor-mediated endocytosis as top pathways. Gene ontology analysis identifying **(A)** molecular function **(B)** cellular component and **(C)** biological process associated with the conserved GPCR interacting proteins. **(D)** KEGG enrichment pathway analysis using reported molecular interactions and relation networks. Size represents the number of conserved proteins that have reported roles in the indicated function, compartment, process, or pathway. Color reflects the extent of q-value enrichment.

To further analyze the GPCR interacting proteins, we utilized the Search Tool for Retrieval of Interacting Genes/Proteins (STRING) database (*61*). Specifically, this database integrates publicly available sources of protein-protein interactions with computational methods to develop a comprehensive protein interaction network. We generated networks of the functional association of our identified hits using STRING, which incorporates both known and predicted interactions **(Figure 4)**. This analysis demonstrates associations that are meant to be specific and meaningful, such as proteins which jointly contribute to a shared function but does not necessarily mean they are physically interacting with each other. We first analyzed protein-protein interactions on the 47 significant hits found in at least two of the four datasets using strict and relaxed thresholds (false discovery rate of 0.1 versus 0.2) (**Figure 4A and 4B)**. The networks which demonstrated the highest combined interaction score had many protein-protein interactions implicated in endocytosis and receptor trafficking like AP2, epsin, and phosphatidylinositol binding clathrin assembly protein (PICALM) (*62*). We then generated networks restricted to only using the top twenty proteins with the largest magnitude in fold change (**Figure 4C and 4D**). The interaction networks are very similar when using both strict and relaxed thresholds. Together, these findings suggest that the proteins which have the largest magnitude in fold change are interacting in a biologically meaningful way based on previously published experiments and databases, but also computationally predicted interactions. In agreement with our previous GO and KEGG analyses, these data further visually represent that these conserved hits are known and predicted to interact with each other, most likely, in receptor internalization and trafficking.

**Figure 4:**
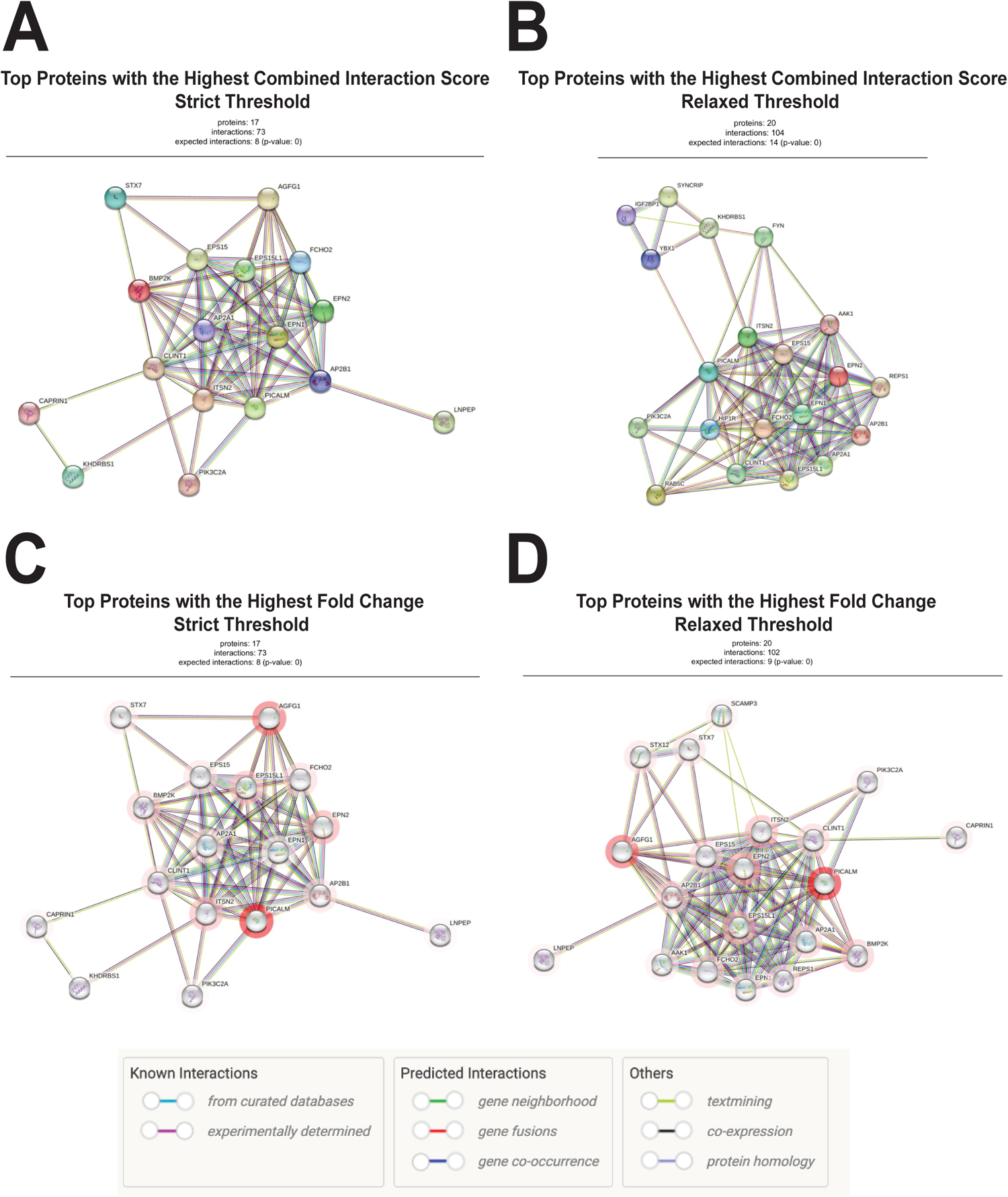
STRING analyses for GPCR interacting proteins. Interaction networks were generated from the STRING database of regulated pathways and using various parameters. Protein-protein interaction network generated using 47 conserved hits at a false discovery rate (FDR) of **(A)** 0.1 and **(B)** 0.2. Shown are only the top interaction networks generated from each report. Protein-protein interaction network generated using only the 20 conserved hits with the largest fold change at an FDR of **(C)** 0.1 and **(D)** 0.2. Magnitude of fold change for each protein is represented by the opacity of the red circle around each node, with a darker red hue representing a larger positive fold change. The basis for the listed interactions between nodes is outlined on the figure insert.

### BMP2K interacts with ACKR3 in the absence of G proteins, β-arrestins, or GRKs

Our proximity labeling experiment and subsequent bioinformatics analyses revealed that bone morphology protein 2-inducible kinase (BMP2K) is significantly upregulated following ACKR3 activation, even in the absence of β-arrestin. Moreover, BMP2K showed significant proximity to GPCRs in the three other datasets as well, suggesting that it may be a conserved GIP. However, to our knowledge, BMP2K has not been studied in the context of GPCR signaling. BMP2K has previously been linked to AP2 phosphorylation and contains a kinase domain that shares a 75% structural similarity to AP2 Associated Kinase 1 (AAK1) (*63–65*). Using a NanoBiT complementation assay, we measured levels of BMP2K recruitment to ACKR3 in WT HEK293T cells, β-arrestin-1/2 KO cells, and GRK-2/3/5/6 KO cells (**Figure 5A**). We found that BMP2K recruited to ACKR3 both in the absence of β-arrestins and GRKs, and that rescue of β-arrestin 1 or 2 had minimal effect on the overall recruitment of BMP2K to ACKR3 (**Figure 5B-5D, Supplemental Figure 3A-3B**).

**Figure 5:**
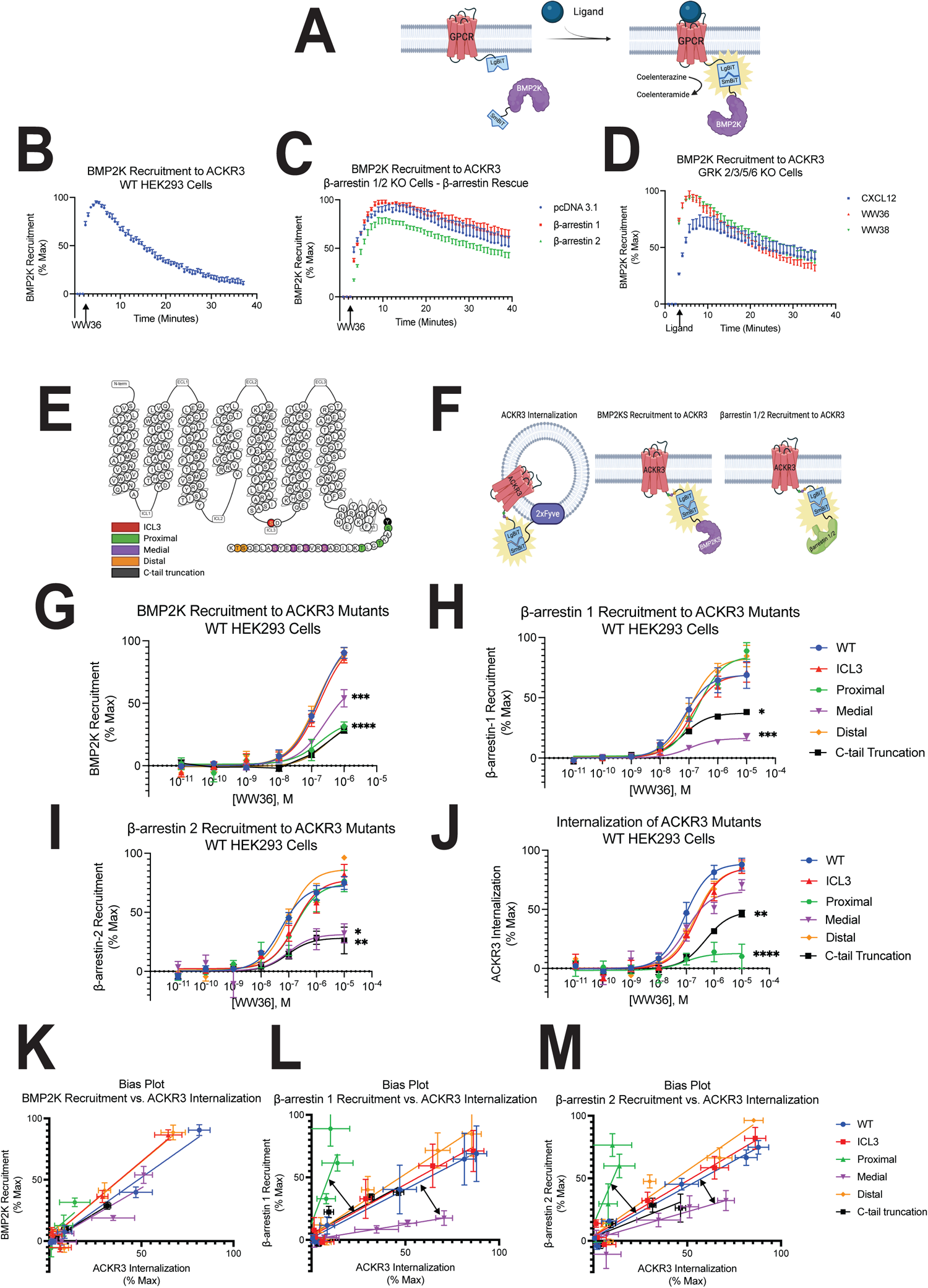
BMP2K and *β*-arrestin recruitment to ACKR3 are dependent on distinct phosphorylation sites. (**A**) Schematic representation of NanoBiT complementation assay used to detect BMP2K recruitment to ACKR3. BMP2K recruitment to ACKR3 in **(B)** WT HEK293 **(C)** β-arrestin 1/2 KO and **(D)** GRK 2/3/5/6 KO cells. Data shown are the mean ± SEM of n=5, n=3, and n=2 respectively. **(E)** Diagram of ACKR3 that shows the specific amino acids mutated to an alanine in each phosphorylation deficient mutant constructed. Black represents a complete C-terminal tail truncation mutant. **(F)** Schematic representation of NanoBiT complementation assays used to measure ACKR3 internalization, BMP2K recruitment to ACKR3, and β-arrestin 1 or 2 recruitment to ACKR3 in HEK293 cells. Agonist concentration-response at five minutes of **(G)** BMP2K **(H)** β-arrestin 1 and **(I)** β-arrestin 2 recruitment to ACKR3, and **(J)** ACKR3 trafficking to early endosomes in HEK293 cells. β-arrestin 1 and 2 recruitment to ACKR3 reflects the mean ± SEM of n=4, and BMP2K recruitment to ACKR3, and ACKR3 trafficking to early endosomes reflects the mean ± SEM of n=5. Significance testing was performed using one-way ANOVA with Dunnet’s post hoc testing comparing luminescent signal at maximum dose between the ACKR3 mutants (WT vs. mutant). Concentration-response curves are normalized to maximum signal observed across all receptors. *P<0.05, **P<0.005, ***P<0.0005, ****P<0.0001. Bias plots demonstrating levels of ACKR3 internalization relative to **(K)** BMP2K **(L)** β-arrestin 1 and **(M)** β-arrestin 2 recruitment to ACKR3 for each of the phosphorylation mutants. Arrows highlight the deviation in best fit lines for the proximal and medial mutants compared to WT in β-arrestin 1 and 2 recruitment level versus ACKR3 internalization.

### Putative ACKR3 phosphorylation sites important for β-arrestins recruitment are distinct from those for ACKR3 internalization and BMP2K interaction

Prior work has shown that specific phosphorylation sites on a GPCR can be important for distinct signaling events, such as G protein activation, and β-arrestin recruitment and conformation (*66–68*). To determine if specific phosphorylation sites on ACKR3 are necessary for BMP2K recruitment, we generated phosphorylation-deficient receptor mutants by mutating putative serine and threonine phosphorylation sites on the receptor’s C-terminus and intracellular loop 3 (ICL3) to alanine (**Figure 5E**). The phosphorylation deficient receptor mutants ranged in targeting ICL3 (ACKR3-S243A), proximal (ACKR3-S335A, T338A, T341A), medial (S347A, S350A, T352A, S355A), and distal (ACKR3-S360A, T361A) phosphorylation sites, as well as a truncation mutant (Y334X) that eliminated the entire C-terminal receptor tail. Alongside BMP2K recruitment, we also measured β-arrestin recruitment and receptor internalization for each of these ACKR3 mutants to evaluate if different phosphorylation sites regulate distinct ACKR3 signaling events (**Figure 5F**).

We found that BMP2K recruitment to ACKR3 depends on the phosphorylation sites mutated. Specifically, the proximal and truncation mutants demonstrate the greatest reduction in BMP2K recruitment, followed by the medial mutant (**Figure 5G**). The recruitment profile of β-arrestins 1 and 2 to the ACKR3 mutants were very similar to one another, with both β-arrestins demonstrating reduced recruitment to the truncation and the medial mutants, while the other ACKR3 mutants were similar to WT ACKR3 (**Figure 5H-5I**). By contrast, phosphorylation sites at the proximal region of the receptor tail were critical for promoting receptor internalization, paralleling the pattern observed for BMP2K recruitment to these mutants (**Figure 5J**). Kinetic tracings for each ACKR3 mutant experiment were also obtained (**Supplemental Figure 3C-3F**).

To qualitatively evaluate the relationships between ACKR3 internalization and β-arrestin or BMP2K recruitment across the ACKR3 mutants, we generated bias plots which enable comparisons of the relative activity between two assays (**Figure 5K-5M**) (*69*). While the ACKR3 mutants exhibit a strong linear correlation between ACKR3 internalization and BMP2K recruitment, β-arrestin 1 and 2 demonstrate deviations in levels of ACKR3 internalization compared to β-arrestin recruitment for the proximal and medial mutants (**Figure 5L-5M)**. The proximal mutant achieves high levels of β-arrestin recruitment despite low levels of ACKR3 internalization, and the medial mutant presents high levels of ACKR3 internalization despite having the least β-arrestin recruitment. These data suggest that specific structural elements regulate ACKR3 signaling events and effector coupling, consistent with existing evidence suggesting that receptor phosphorylation sites can promote distinct canonical and noncanonical signaling events (*66*). Importantly, these data demonstrate that BMP2K interacts with ACKR3 in a manner that is independent from β-arrestins and may have a functional relevance to receptor internalization.

### BMP2K interacts with many therapeutically relevant GPCRs with varying dependence on β-arrestin expression

The finding that BMP2K recruits to ACKR3 independent of β-arrestins (and G proteins) prompted further inquiry into whether BMP2K recruits to other GPCRs, and if its interaction with other receptors is similarly β-arrestin-independent. We selected four well-studied and therapeutically-relevant GPCRs with varying physiological functions and Gα subunit activation: AT_1_R, V_2_ vasopressin receptor (V_2_R), β_2_AR, and C-X-C motif chemokine receptor 3 (CXCR3). Using the same NanoBiT complementation assay as in Figure 5, we measured BMP2K recruitment to these receptors in β-arrestin 1/2 KO cells with and without β-arrestin 1 or 2 rescue.

These four GPCRs demonstrated unique patterns of BMP2K recruitment that were largely dependent upon the presence of β-arrestin. At the AT_1_R, β-arrestin 1 and 2 each promoted the recruitment of BMP2K to the receptor, but little BMP2K recruitment was detected when both isoforms were absent (**Figure 6A**). BMP2K recruitment to V_2_R and β_2_AR similarly required β-arrestins, with β-arrestin 1 inducing greater BMP2K recruitment than β-arrestin 2 upon stimulation of each receptor (**Figure 6B-C**). By contrast, in the absence of β-arrestins, BMP2K recruited to CXCR3 slowly (**Figure 6D**). This recruitment pattern was expedited following addback of β-arrestin 1, but eliminated with the addback of β-arrestin 2. These findings indicate that BMP2K interacts with various GPCRs, but that the specific mechanism and temporal pattern of this interaction is largely receptor- and β-arrestin dependent.

**Figure 6:**
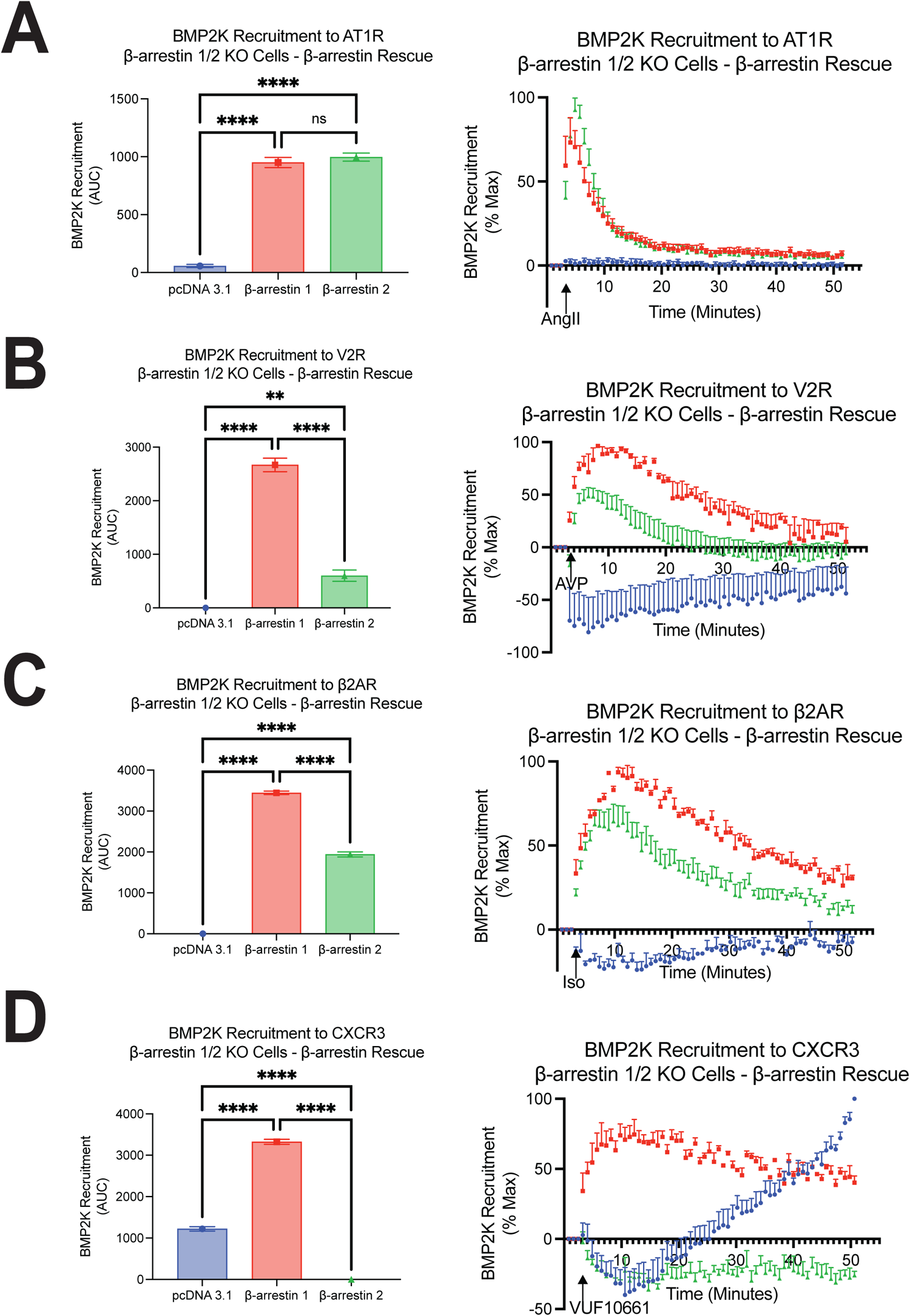
BMP2K interactions with different GPCRs has different requirements for *β*-arrestins. Area under the curve and kinetic time-course of BMP2K recruitment to **(A)** AT_1_R following stimulation with 10µM angiotensin II, **(B)** V_2_R following stimulation with 10µM arginine vasopressin, **(C)** β_2_AR following stimulation with 10µM isoproterenol, and **(D)** CXCR3 following stimulation with 10µM VUF10661. Data shown are the mean AUC ± SEM and mean ± SEM of between n=3-5. Significance testing was performed using one-way ANOVA with Turkey’s post hoc testing comparing mean AUC between treatment groups (pcDNA vs. β-arrestin 1 vs. β-arrestin 2). Kinetic time-course is normalized to maximum signal observed across all treatment groups. *P<0.05, **P<0.005, ***P<0.0005, ****P<0.0001.

## DISCUSSION

Noncanonical GPCR signaling is an emerging area of study that adds to our understanding of the complexity of GPCR signaling and how a single agonist:receptor interaction results in the activation of complex cellular and physiologic processes. For instance, recent findings suggest that Gαi can form a complex with β-arrestin and scaffold MAPK, even at receptors that do not canonically couple to Gαi (*23, 70*). There have also been reported ternary complexes between G proteins, β-arrestins, and GPCRs that can modulate GPCR signaling (*22, 71, 72*). However, G proteins and β-arrestins are not the only modulators of GPCR signaling, with evidence that GIPs, such as PDZ domain-containing proteins and A-Kinase Anchoring Proteins (AKAPs), independently regulate GPCR signaling (*3, 73*). Because of recent evidence demonstrating that ACKR3 can signal without G proteins or β-arrestins, we hypothesized that this receptor could serve as a model GPCR to further explore noncanonical interacting partners of GPCR signaling.

We found that ACKR3 internalization is preserved in both β-arrestin 1/2 KO and GRK 2/3/5/6 KO cell lines. Our data, alongside prior work which highlight the importance of receptor phosphorylation to achieve ACKR3 function, suggest that necessary ACKR3 phosphorylation can be GRK-independent, and may involve either basal phosphorylation and/or phosphorylation mediated by other kinases (*37, 44*). We then used proximity labeling to identify noncanonical signaling effectors of ACKR3, a receptor that does not couple to G proteins, in a cell line devoid of β-arrestin. We tracked ACKR3:protein proximity using APEX2 labeling, and identified ∼100 proteins with significantly changed proximity to ACKR3 following receptor stimulation.

Most of these proteins have recorded involvement in various forms of the endocytic machinery, such as cargo transport to the Golgi, and vesicle recycling to the plasma membrane. VPS29, which demonstrated the largest expression fold change, is not only a protein component of the endosomal cargo recycling machinery called the retromer complex, but has also been implicated in a newly discovered mechanism of endosomal recycling called the “retriever” complex (*52*). The retromer complex carries endosomal cargo to the trans-golgi network and the plasma membrane, and has been implicated at the parathyroid hormone receptor (PTHR) (*74–76*). The retriever complex is part of a larger 15 protein assembly called the “commander” complex, which participates in endosomal trafficking between the plasma membrane, trans-golgi network, nucleus, and lysosomes (*52, 77*). While it cannot be directly deduced whether the VPS29 identified in our APEX experiment is representative of one or both the retromer and retriever complex in ACKR3 trafficking, it is possible that the retriever complex, and by extension the commander complex, may be involved in GPCR endosomal sorting and trafficking as well (*78*). Another identified protein, the AKT interacting protein (AKTIP), is a component of the FHF complex, which is involved in attaching cargo to dynein for retrograde transport (*79, 80*). These findings suggest that numerous protein complexes with roles in endocytic trafficking, such as the retromer, commander, and FHF complexes, may regulate ACKR3 intracellular trafficking independently of β-arrestins.

Our data reflects what is, to our knowledge, the first implementation of proximity labeling to study the interaction profile of a GPCR signaling without G protein or β-arrestin mediators. By comparing the proteins identified in the current dataset to three similar experiments at other conventional GPCRs, we identified 17 proteins with significant GPCR interactions conserved across all datasets. Further pathway analyses of these proteins using GO, KEGG, and STRING revealed clathrin-mediated endocytosis as a highly conserved mechanism across each of these GPCRs. This is likely due to AP2’s ability to bind both β-arrestins and various motifs found in some GPCRs, thus facilitating β-arrestin dependent and independent clathrin-mediated endocytosis (*27*). Many of the protein effectors implicated in endosomal sorting and recycling that were detected in our dataset were not identified in the others. This finding may be due to differences in experimental conditions, dynamics of ACKR3 endocytosis, or the result of ACKR3 employing different endosomal machinery than other GPCRs.

These analyses allowed us to identify unique β-arrestin-independent, ACKR3-interacting proteins which appear to function as conserved GPCR effectors. We identified BMP2K as one of the top hits in each proximity labeling dataset, and found that BMP2K recruits to ACKR3 in a β-arrestin- and GRK-independent fashion. We developed ACKR3 phosphorylation site deficient mutants to show that there are specific phosphorylation sites on the C-terminus of ACKR3 involved in BMP2K recruitment, with proximal sites demonstrating the most pronounced effect. In contrast, β-arrestin recruitment patterns were most influenced by medial sites, as reflected in earlier studies (*44*). Consistent with our finding that β-arrestin is not required for BMP2K recruitment, we discovered that the putative phosphorylation sites which regulate BMP2K recruitment to ACKR3 are different from those which regulate β-arrestin recruitment. Together, these data further suggests that BMP2K is a novel interacting partner at ACKR3, and that it likely possesses a receptor binding interface that is distinct from β-arrestin. Of note, BMP2K recruitment to the phosphorylation-deficient ACKR3 mutants was strikingly similar to the pattern of receptor internalization for these mutants, suggesting a relationship between BMP2K and receptor endocytosis. It is possible that BMP2K may induce phosphorylation of ACKR3, thereby promoting receptor internalization, or that ACKR3 internalization may promote BMP2K recruitment. Further studies are needed to elucidate the temporal ordering and significance of BMP2K recruitment in ACKR3 internalization.

Given our discovery of a novel G protein and β-arrestin independent effector of ACKR3, we also studied BMP2K engagement at four other GPCRs. Interestingly, we found that the kinetics and dependence on β-arrestin for BMP2K:GPCR interaction patterns varied across receptors. While ACKR3 recruits BMP2K without β-arrestins, other GPCRs had specific β-arrestin isoform requirements to promote BMP2K recruitment. While the AT_1_R exhibited a similar increase in recruitment of BMP2K following rescue with β-arrestin 1 or 2, only β-arrestin 1 rescue significantly increased BMP2K recruitment to CXCR3. These findings suggest that transducers like β-arrestin may be necessary to promote BMP2K engagement at some receptors but may be dispensable at others. It is well established that β-arrestin can act as a scaffold protein, binding to and promoting cargo proteins like clathrin and AP2 to the receptor (*81*). With BMP2K’s reported role in phosphorylating AP2, a necessary step in clathrin-mediated endocytosis, it is possible that β-arrestin may scaffold BMP2K to certain GPCRs that require β-arrestin for their function (*63*).

Our experiments and analyses led to the identification of several protein hits conserved across GPCRs. However, it is important to note that these results are limited in their physiological relevance due to our experimental design. ACKR3 is not endogenously expressed in HEK293 cells, which limits the ability to associate our findings with ACKR3’s known role in inflammation, cardiac development, and neuronal migration (*29, 82–84*). Further studies of ACKR3’s function would likely benefit from investigating the receptor in cell types where endogenous ACKR3 signaling and function have been recorded, such as in macrophages and cardiomyocytes (*85, 86*). While CXCL12 is the endogenous ligand of ACKR3, it can also activate the C-X-C motif chemokine receptor 4 (CXCR4), known to be expressed in HEK293 cells (*49*). Because ACKR3 can dimerize with CXCR4, some of the protein interactions identified may be CXCR4-dependent (*28*). However, our experimental design requires a protein has to be within a 20-nanometer distance from the APEX2 enzyme in order to be biotinylated, resulting in minimal chances of off-target protein interactions (*47*). Importantly, CXCR4 was not identified as a protein hit in our dataset, further suggesting that the interactome identified in this manuscript is that of ACKR3 alone.

Given the emphasis in canonical GPCR signaling on G proteins and β-arrestins, efforts to develop “biased” GPCR therapeutics that preferentially activate one pathway over the other have been extensively pursued (*3, 87, 88*). Despite these efforts, only one biased GPCR therapeutic has received Food and Drug Administration (FDA) approval and has demonstrated limited efficacy compared to other available treatment options (*89*). One reason for this lack of anticipated therapeutic benefit could be that associating far downstream cellular outcomes directly with levels of G protein versus β-arrestin signaling fails to recognize the influence of other factors that impact GPCR functionality (*90*). Considering the roles of G proteins and β-arrestins independently of one another fails to recognize not only how the two influence one another, but also that there is a diverse array of other effector proteins known to directly interact with and modify GPCRs (*17, 19*). G proteins and β-arrestins are also not the only modulators of GPCR signaling, and with emerging evidence showing the ability of GPCRs to signal in the absence of both, one should consider the roles of GIPs more thoroughly. Implementing the use of a systems pharmacology approach to thoroughly investigate the functional consequences of these different GIPs on signaling outcome may lead to a more holistic understanding of GPCR signaling, which could result in the development of more specific and efficacious therapeutics.

To our knowledge, this data serves as the first known evidence of BMP2K’s involvement in GPCR signaling, yet its functional mechanism has yet to be elucidated. Moreover, we have identified many new conserved protein effectors which may modulate GPCR signaling in ways that have yet to be explored. Given the nature of our experimental design, we anticipate that these other effectors will also demonstrate pleiotropic signaling behavior at different GPCRs. These findings highlight the potential behind exploring noncanonical signaling proteins, both in its capacity for revealing novel ways to modulate GPCR signaling, and in its expansion of potential drug targets for future therapeutics that may lead to more specified treatment responses. Although GPCR signaling is primarily mediated by G proteins and β-arrestin, we demonstrate that there are likely many other effectors which contribute to generating highly complex and specific cellular and physiologic responses.

## ACKNOWLEDGEMENTS

We thank the members of the Duke Proteomics and Metabolomics Core Facility who managed the collection and analysis of the ACKR3-APEX2 mass spectrometry data: Tricia Ho (sample preparation, data collection, data analysis, report writing), Greg Waitt (data analysis), and Erik J. Soderblom (study design, data analysis, report writing, scientific oversight); Kevin Zheng for his thoughtful perspective on APEX dataset statistical comparisons; Gayathri Viswanathan and Nour Nazo for valuable laboratory insight and assistance. Graphical figures were created using BioRender.

## FUNDING

National Institute of General Medical Sciences T32GM007171 (D.S.E)

National Institute of General Medical Sciences R01GM122798 (S.R.)

National Institutes of Health T32GM148377 (J.G)

Duke University Medical Scientist Training Program (D.S.E)

University of Pennsylvania Medical Scientist Training Program (J.G)

American Heart Association Predoctoral Fellowship 20PRE35120592 (D.S.E.)

American Heart Association Predoctoral Fellowship 23PRE1019796 (U.P)

National Institutes of Health Intramural Research Training Award Fellowship (C.H.)

Duke Cardiovascular Research Center Undergraduate Research Experiences Fellowship (C.H.)

## AUTHOR CONTRIBUTIONS

Conceptualization, C.H., J.G., D.S.E., S.R.

Methodology, C.H., J.G., D.S.E., N.D.C., S.R.

Investigation, C.H., J.G., D.S.E., N.D.C., U.P., S.D., H.R., A.C.

Writing — Original Draft, C.H., J.G., D.S.E., N.D.C.

Writing – Reviewing & Editing, C.H., J.G., D.S.E., N.D.C., U.P., S.D., H.R., A.C., S.R.

Visualization, C.H., J.G., D.S.E., N.D.C., S.R.

Supervision and Funding Acquisition, S.R.

## DATA AND MATERIALS AVAILABILITY

### Lead Contact

Further information and requests for resources and reagents should be directed to and will be fulfilled by the lead contact, Sudarshan Rajagopal.

### Materials Availability

All plasmids generated in this study will be distributed upon request.

### Data and Code Availability

All data reported in this paper will be shared by the lead contact upon request.

## MATERIALS AND METHODS

### Materials Availability

All plasmids generated in this study will be distributed upon reasonable request.

### Data and Code Availability

- The mass spectrometry proteomics data have been deposited to the ProteomeXchange Consortium via the PRIDE partner repository with the dataset identifier PXD047836 (*91*)
- This paper does not report original code.
- All data reported in this paper and any additional information required to reanalyze the data reported in this paper is available from the lead contact upon request.

## EXPERIMENTAL MODEL AND SUBJECT DETAILS

### Bacterial Strains

XL-10 Gold ultracompetent E. coli (Agilent) were used to express all constructs used in this manuscript.

### Cell Lines

Human Embryonic Kidney (HEK293) cells were grown in minimum essential media (MEM) supplemented with 10% fetal bovine serum (FBS) and 1% penicillin/streptomycin at 37°C and 5% CO2. Δβ-arrestin1/2, and ΔGRK2/3/5/6 CRISPR/Cas9 KO HEK293 cells were provided by Asuka Inoue, Tohoku University, Japan, and validated in prior publications (*48*).

## METHOD DETAILS

### Generation of Constructs

Construct cloning was performed using conventional techniques including restriction digest and overlap extension PCR cloning (New England Biolabs), and site-directed mutagenesis (QuikChange Lightning kit, Agilent Technologies). Linkers separating fluorescent proteins or luciferases and the cDNA for receptors, location markers, or other proteins were flexible and ranged between 6 and 16 amino acids. The position of the fluorescent proteins or luciferases relative to the construct of interest is indicated by its positioning either before or after the construct name. We utilized the 2x-Fyve sequence (RKHHCRACG) from the hepatocyte growth factor-regulated tyrosine kinase substrate to detect ACKR3 recruitment to early endosomes, and the CAAX membrane sequence to detect β-arrestin recruitment to the plasma membrane. All BMP2K constructs were derived from a pcDNA3-BMP2K plasmid provided by Jaroslaw Cendrowski, International Institute of Molecular and Cell Biology, Warsaw (*92*). All BMP2K experiments were performed using the human short splicing variant of BMP2K. The ACKR3-APEX2 plasmid was engineered using the backbone of an AT1R-APEX2 construct provided by Howard Rockman, Duke University, Durham (*39*). Mouse β-arrestin isoforms 1 and 2 were used for experiments involving β-arrestin 1 or 2 rescue and all experiments measuring β-arrestin recruitment to the plasma membrane and to GPCRs. Bovine GRK2, human GRK3 Isoform 1, human GRK5, and human GRK6 isoform B were used for experiments involving GRK rescue in the GRK2/3/5/6 KO cell line. Human ACKR3, AT_1_R, β_2_AR, CXCR3A, and V_2_R isoform 1 were used in this study.

### Cell Culture and Transfection

For both luminescence-based assays (NanoBiT split) and BRET based assays, cells were transiently transfected using polyethylenimine (PEI). This transfection method involves replacing the cell culture media 30 minutes prior to transfection. Following the re-media of the cells, plasmid constructs were prepared and suspended in Opti-MEM (Gibco) which was added to reach a final volume of 100μL. In another tube, a 1 mg/mL concentration of PEI was added to Opti-MEM to reach a final volume of 100μL. For the experiments shown in this paper, the PEI:DNA ratio was 3:1, meaning that for every 1μg of DNA plasmid added, 3μL of PEI was also added. After allowing the PEI tube to incubate for 5 minutes, the 100μL PEI solution was added to the 100μL DNA solution and gently pipetted. After allowing the combined solution to incubate to room temperature for 10-15 minutes, the 200μL solution was added to one well of a 6-well plate.

### BRET and NanoBiT Luciferase Assays

For the HEK293, β-arrestin 1/2 KO, and GRK 2/3/5/6 KO cells used, cells were seeded in 6 well plates averaging approximately 750,000 cells per well. For all BRET and NanoBiT assays performed, the cells were transiently transfected with their corresponding constructs using the aforementioned PEI method. ACKR3 internalization in the HEK293, β-arrestin ½ KO, and GRK 2/3/5/6 KO cell lines were measured using a BRET complementation assay that included a C-terminal ACKR3-Renilla Luciferase II (RLucII) and 2xFyve-mVenus to measure ACKR3 recruitment to early endosomes.

β-arrestin recruitment to the plasma membrane in parental and ACKR3-APEX2 expressing β-arrestin 1/2 KO cells was assessed using a NanoBiT complementation assay involving SmBiT-β-arrestin-2 and LgBiT-CAAX. BMP2K recruitment was similarly assessed using a NanoBiT complementation assay where, for studying recruitment of BMP2K to various GPCRs, SmBiT-BMP2K was tagged and transfected in with either ACKR3-LgBiT, AT_1_R-LgBiT, β_2_AR-LgBiT, CXCR3-LgBiT, or V_2_R-LgBiT. In constructing the phosphodeficient mutants of ACKR3, each of the ACKR3 mutants was cloned using ACKR3-LgBiT as the template plasmid. Internalization with the receptor mutants was measured using the early endosomal marker 2xFyve-SmBiT. C-terminal BMP2K-SmBiT was used to measure BMP2K recruitment to the ACKR3 mutant receptors, and SmBiT-β-arrestin-1 and 2 were transfected alongside the ACKR3 mutants to measure β-arrestin recruitment to the ACKR3 mutant receptors.

Twenty-four hours following transfection, cells were washed with phosphate buffered saline (PBS) and collected using trypsin. Cells were resuspended in clear minimum essential medium (Gibco) with the addition of 2% FBS, 1% P/S, 10 mM HEPES, 1x GlutaMax (Gibco), and 1x Antibiotic-Antimycotic (Gibco). Upon resuspension, cells were plated into clear or white-bottomed, white-walled Costar 96-well plates with each well averaging approximately 50,000 to 100,000 cells.

The following day, the media was removed and replaced with 80μL of Hanks’ Balanced Salt Solution (HBSS) supplemented with 20mM HEPES and 3µM coelenterazine h for all BRET and NanoBiT complementation assays (Nanolight Technology, Pinetop, AZ), after which the cells were incubated at 37°C for 10 to 15 minutes. For all BRET assays, a standard 480nm RLuc emission filter and a 530nm long pass filter was used to quantify the net BRET ratio (Chroma Technology Co., Bellows Falls, VT).

For NanoBiT complementation and BRET experiments, three initial reads were taken prior to stimulating the cells with ligand to quantify a baseline luminescence or BRET ratio before adding ligand. Cells were stimulated with either a vehicle control (HBSS with 20 mM HEPES) or with the indicated ligand concentration. The change in luminescence or BRET ratio following ligand stimulation was normalized to vehicle treatment. The BRET experiment was quantified using a BRET ratio, where the acceptor signal (mVenus) was divided by the luminescent signal (RlucII). A net BRET ratio was quantified by normalizing to the BRET ratio of the vehicle treatment. For NanoBiT complementation assays, changes in luminescence were normalized to the maximum luminescence change. The 96 well plates were read using a BioTek Synergy Neo2 plate reader or a Berthold Mithras LB 940 set at 37°C. All readings were performed using a kinetic protocol.

### Confocal microscopy

ACKR3-APEX2 β-arrestin ½ KO Cells were plated in rat-tail-collagen-coated 35 mm glass bottomed dishes (MatTek Corporation, Ashland, MA). Cells were imaged with a Zeiss CSU-X1 spinning disk confocal microscope using the corresponding lasers to excite GFP (480 nm). Confocal images were arranged and analyzed using ImageJ (NIH, Bethesda, MD).

### Generation of Lentivirus and stably expressing ACKR3-APEX2 **β**-arrestin 1/2 KO cell line

After generating the aforementioned ACKR3-APEX2 fusion construct, we generated lentivirus as previously described using a second generation approach (*66*). Briefly, the receptor construct was cloned into a pLenti plasmid backbone consisting of the receptor underneath a CMV promoter. HEK293 cells were transfected using a calcium phosphate method with the pLenti receptor containing plasmid, envelope vector (pMD2.G), and packaging vector (psPAX2). 16h post-transfection, the HEK293 cell media was changed. At 40h, the media was harvested, filtered, and replaced. At 64h post transfection, the viral containing media was harvested and added to the aliquot obtained at 40h, and virus was concentrated using the Lenti-X concentrator (Takara Bio, Japan) and viral titer was determined using qPCR per the manufacturer guidelines (ABM, Canada). β-arrestin 1/2 KO cells were transduced with virus multiplicity of infection of 80–100 in the presence of polybrene at 8mg/mL and left to incubated for 72h. Cells were then incubated with puromycin to select only for cells that were successfully transduced with lentivirus.

### Fluorescence-Activated Cell Sorting (FACS)

Cells that were successfully transduced with lentivirus and underwent puromycin selection were then sorted via FACS to obtain cells that express receptor to a similar degree. Specifically, cells were harvested and washed with PBS containing 2.5% FBS and 1.5mM EDTA. Allophycocyanin (APC) conjugated Anti-Human CXCR7 Monoclonal Antibody (Biolegend, San Diego, CA) was added to the cells per the manufacturers recommendation and allowed to incubate for 25 minutes on ice in the dark. Cells were then washed with PBS with 2.5% FBS and 1.5mM EDTA and resuspended in PBS containing DNAse (10µg/mL). Cells were filtered through a sterile 30µm filter and sorted using FACS on an Astrios (Beckman Coulter) sorter at the Duke University Cancer Institute. Analyses were conducted using FlowJo version 10 software.

### APEX2 proximity labeling

We followed an APEX2 protocol as described in prior publications (*47*). Stably expressing ACKR3-APEX2 β-arrestin 1/2 KO cells were placed in a 6cm dish with starvation media (MEM, 1% P/S, and 0.5% FBS) for at least four hours prior to experimentation. 30 minutes to 1 hour prior to labeling, biotin phenol (Iris Biotech) in DMSO stored at −80°C was added to the cells to a final concentration of 500µM. Cells were stimulated for a total of 3 minutes with CXCL12 at a final concentration of 100nM. During the final minute of agonist simulation, a solution of hydrogen peroxide in PBS was added to a final concentration of 1mM. Next, the cell media containing biotin phenol, CXCL12, and hydrogen peroxide was aspirated and cells were washed with 3mL of fresh quenching buffer (10mM sodium ascorbate, 5mM Trolox, 10mM sodium azide) three times to stop the proximity labeling. Cells were then lysed using 500mL of radioimmunoprecipitation (RIPA) buffer (150mM NaCl, 1% Nonidet P-40, 0.5% sodium deoxycholate 0.1% sodium dodecyl sulfate, 50mM Tris pH 7.4) supplemented with cOmplete protease inhibitor (Sigma-Aldrich) and quenching reagents (10mM sodium azide, 10mM sodium ascorbate, and 5mM Trolox). Cells were collected, left to sit on ice for two minutes, and then rotated at 4°C for 1 hour, followed by subsequent centrifugation at 17000G for 10 minutes where the supernatant was then collected and stored for further analyses.

### Immunoprecipitation of Biotinylated Proteins

Isolation of biotinylated proteins was performed using streptavidin magnetic beads as previously described (*47*). Streptavidin magnetic beads were washed twice with 1mL of RIPA lysis buffer. Whole cell lysate samples were incubated and rotated with 30µL of streptavidin magnetic beads and 500µL of RIPA buffer for 1 hour at room temperature. The beads were then collected using a magnetic rack, and the supernatant was removed and stored for later immunoblotting analysis. The remaining magnetic beads were washed twice with 1mL of RIPA lysis buffer, once with 1mL of 1M KCl, once with 1mL of 0.1M sodium carbonate, once with 1mL of 2M urea in 10mM Tris-HCl (pH 8.0), and then twice again with 1mL of RIPA lysis buffer. To elute biotinylated proteins from the magnetic beads, the samples were boiled for 10 minutes in 100µL of 25mM Tris, 50mM NaCl, 10mM DTT, 2% SDS, and 5mM Biotin. We then vortexed the beads briefly, and cooled the sample on ice. The samples were then placed on the magnetic rack and the eluate was collected and placed on ice.

### Immunoblotting and Coomassie Stain

Experiments were conducted as previously described (*93*). Lysates or eluents were resolved on SDS-10% polyacrylamide gels, transferred to nitrocellulose membranes, and immunoblotted with the indicated primary antibody overnight at 4°C. Specifically, we used A1CT (courtesy of the Lefkowitz Laboratory (*94*)) to detect expression of β-arrestin1/2 and anti-alpha-tubulin antibodies (ThermoFisher) to detect expression of tubulin.

Peroxidase–conjugated polyclonal donkey anti-rabbit immunoglobulin (IgG) or polyclonal sheep anti-mouse IgG were used as secondary antibodies. A peroxidase-conjugated streptavidin was used to detect biotinylated proteins. Immune complexes on nitrocellulose membrane were imaged as previously described by incubating cells in a mixture of a solution of 100mM Tris (pH 8.6) and 1.25mM sodium luminol in water with a solution of 1.1mg/mL para-hydroxy coumaric acid in DMSO (*95*). Cells were imaged using a ChemiDoc MP Imaging System (Bio-Rad). Representative immunoblots demonstrate all samples run, but are cropped to highlight the bands of interest. Coomassie staining was performed on polyacrylamide gels by staining the gels in a Coomassie solution (50% methanol, 10% acetic acid, 39.75% deionized water, and 0.25% Coomassie Blue R-250) overnight. Gels were then destained for 2-4 hours in 5% methanol, 7.5% acetic acid, and 87.5% deionized water and then imaged.

### Mass Spectrometry (MS) Sample Preparation

Mass spectrometry was performed by the Duke Proteomics and Metabolomics Core Facility (Durham, NC). 6 samples (3 samples treated with vehicle control and 3 samples treated with CXCL12) were kept at −80°C until processing. Samples were spiked with undigested bovine casein at a total of either 120 or 240 pmol as an internal quality control standard. Next, samples were supplemented with 17.6 μL of 20% SDS, reduced with 10 mM dithiolthreitol for 30 min at 80°C, alkylated with 20mM iodoacetamide for 30 min at room temperature, then supplemented with a final concentration of 1.2% phosphoric acid and 1012 μL of S-Trap (Protifi) binding buffer (90% MeOH/100mM TEAB). Proteins were trapped on the S-Trap micro cartridge, digested using 20 ng/μL sequencing grade trypsin (Promega) for 1 hr at 47C, and eluted using 50 mM TEAB, followed by 0.2% FA, and lastly using 50% ACN/0.2% FA. All samples were then lyophilized to dryness. Samples were resolubilized using 120 μL of 1% TFA/2% ACN with 25 fmol/μL yeast ADH.

### LC-MS/MS Analysis

Quantitative LC/MS/MS was performed on 2 μL (16.6% of total sample) using an MClass UPLC system (Waters Corp) coupled to a Thermo Orbitrap Fusion Lumos high resolution accurate mass tandem mass spectrometer (Thermo) equipped with a FAIMSPro device via a nanoelectrospray ionization source. Briefly, the sample was first trapped on a Symmetry C18 20 mm × 180 μm trapping column (5 μl/min at 99.9/0.1 v/v water/acetonitrile), after which the analytical separation was performed using a 1.8 μm Acquity HSS T3 C18 75 μm × 250 mm column (Waters Corp.) with a 90-min linear gradient of 5 to 30% acetonitrile with 0.1% formic acid at a flow rate of 400 nanoliters/minute (nL/min) with a column temperature of 55C. Data collection on the Fusion Lumos mass spectrometer was performed for three difference compensation voltages (−40v, −60v, - 80v). Within each CV, a data-dependent acquisition (DDA) mode of acquisition with a r=120,000 (@ m/z 200) full MS scan from m/z 375 – 1500 with a target AGC value of 4e5 ions was performed. MS/MS scans were acquired in the ion trap in Rapid mode with a target AGC value of 1e4 and max fill time of 35 ms. The total cycle time for each CV was 0.66s, with total cycle times of 2 sec between like full MS scans. A 20s dynamic exclusion was employed to increase depth of coverage. The total analysis cycle time for each injection was approximately 2 hours.

### Quantitative MS Data Processing

Following LC-MS/MS analyses, data were imported into Proteome Discoverer 2.5 (Thermo Scientific Inc.). In addition to quantitative signal extraction, the MS/MS data was searched against the SwissProt *H. sapiens* database (downloaded in Nov 2019) and a common contaminant/spiked protein database (bovine albumin, bovine casein, yeast ADH, etc.), and an equal number of reversed-sequence “decoys” for false discovery rate determination. Sequest with Infernys enabled (v 2.5, Thermo PD) was utilized to produce fragment ion spectra and to perform the database searches. Database search parameters included fixed modification on Cys (carbamidomethyl) and variable modification on Met (oxidation). Search tolerances were 2ppm precursor and 0.8Da product ion with full trypsin enzyme rules. Peptide Validator and Protein FDR Validator nodes in Proteome Discoverer were used to annotate the data at a maximum 1% protein false discovery rate based on q-value calculations. Note that peptide homology was addressed using razor rules in which a peptide matched to multiple different proteins was exclusively assigned to the protein has more identified peptides. Protein homology was addressed by grouping proteins that had the same set of peptides to account for their identification. A master protein within a group was assigned based on % coverage.

Prior to imputation, a filter was applied such that a peptide was removed if it was not measured in at least 2 unique samples (50% of a single group). After that filter, any missing data missing values were imputed using the following rules; 1) if only one single signal was missing within the group of three, an average of the other two values was used or 2) if two out of three signals were missing within the group of three, a randomized intensity within the bottom 2% of the detectable signals was used. To summarize to the protein level, all peptides belonging to the same protein were summed into a single intensity. These protein levels were then subjected to a normalization in which the top and bottom 10 percent of the signals were excluded and the average of the remaining values was used to normalize across all samples. We calculated fold-changes between vehicle and CXCL12 based on the protein expression values and calculated two-tailed heteroscedastic t-test on log2-transformed data for each of these comparisons.

### Quantification and Statistical Analysis

Data were analyzed in Microsoft Excel (Redmond, WA) and graphed in Prism 9.5 (GraphPad, San Diego, CA). Kinetic time course data was acquired by tracking changes in luminescence signal of the highest non-toxic dose (compared to a baseline of no ligand treatment), with the maximum change in luminescence normalized to 100. Dose-response curves were fitted to a log agonist versus stimulus with three parameters (span, baseline, and EC50), with the minimum baseline corrected to zero. Statistical tests were performed using a one-way ANOVA followed by Tukey’s or Dunnet’s multiple comparisons test when comparing treatment conditions. When comparing treatment conditions in time-response assays, a one-way ANOVA of treatment and AUC was conducted. A significant interaction effect was defined as having an adjusted P value less than 0.05 (*P* < 0.05). Further details of statistical analysis and replicates is included in the figure legends. Lines are representative of the mean and error bars indicate the SEM. Independent replicates of plate-based experiments were procured by at least two different investigators when feasible.

### MS Data Postprocessing and Protein Identifier Mapping

Rajagopal, Kruse, Rockman, and Von Zastrow datasets were filtered for treatment time points at 3 minutes, 2 minutes, 1.5 minutes, and 3 minutes respectively, and all fold change data with respect to control or vehicle conditions across datasets were log_2-_transformed. Values above Q3 + 3x interquartile range (IQR) or below Q1 - 3xIQR were considered extreme points and removed. All protein accessions were mapped to the UniProtKB database gene names using the website https://www.uniprot.org/id-mapping for human-readable labeling of accessions as well as for input into the Gene Ontology/KEGG Enrichment analysis. Additionally, these UniProt identifiers were mapped to Entrez Gene identifiers using the *mapIDs* function from “AnnotationDbi” R package and the genome annotation database from the “org.Hs.eg.db” R package for input into the STRINGdb analysis (*59, 96*). Accessions were labeled “conserved” if they were present in all four datasets. Accessions were labeled significant if both the fold change was less than 0.5 or greater than 2 (0.5 < fold change < 2) and had a False Discovery Rate (FDR)-corrected combined Fisher p-value strictly less than 0.1.

### Combined Fisher’s P-value Calculation

Combined p-values were constructed using Fisher’s method on the Rajagopal, Rockman, and Von Zastrow datasets. Briefly, each two-sided p-value was first converted to a one-sided p-value using the *metap* function from the “metap” R package (*97*). Then, the p-values were grouped by accession and combined across datasets using the *sumlog* function from the “metap” R package (*97*). Finally, the resultant combined Fisher p-values were corrected for multiple hypothesis testing to produce the FDR-corrected combined Fisher p-value.

### Gene Ontology/KEGG Enrichment and STRINGdb Protein-Protein Interaction and Clustering

Gene Ontology (GO) Enrichment and KEGG Database Enrichment were performed using the *enrichGO* and *enrichKEGG* functions respectively from the R package “clusterProfiler” (*59, 98*). GO enrichment was assessed for the following categories: molecular function (MF), biological process (BP), cellular compartment (CC), and all three categories at once (ALL). GO and KEGG Enrichment categories were filtered for FDR-corrected p-value < 0.1. STRINGdb (version 11.5, species 9606) protein-protein Interaction, network visualization, and clustering analyses were performed using the *map*, *get_interactions*, *plot_network*, and *cluster* functions from the R package “STRINGdb” (*59, 99*). STRINGdb minimum protein-protein interaction score threshold was set to 200 (or 0.200 on the STRINGdb webpage), and the FastGreedy algorithm was used for STRINGdb clustering with default parameters.

### Waterfall and Volcano Visualizations

Volcano plots were constructed using functions from the “tidyverse” R package on the median log_2_ fold change across datasets for each accession and the associated negative log_10_ of the FDR-corrected Fisher combined p-value with a custom R-script (*100*). The accessions from the Kruse dataset, which did not report p-values, were assigned the FDR-corrected combined Fisher p-value to utilize the Kruse log_2_ fold changes in the volcano plots. Waterfall plots were similarly constructed using the individual log_2_ fold changes of each accession for each dataset.

## KEY RESOURCES TABLE

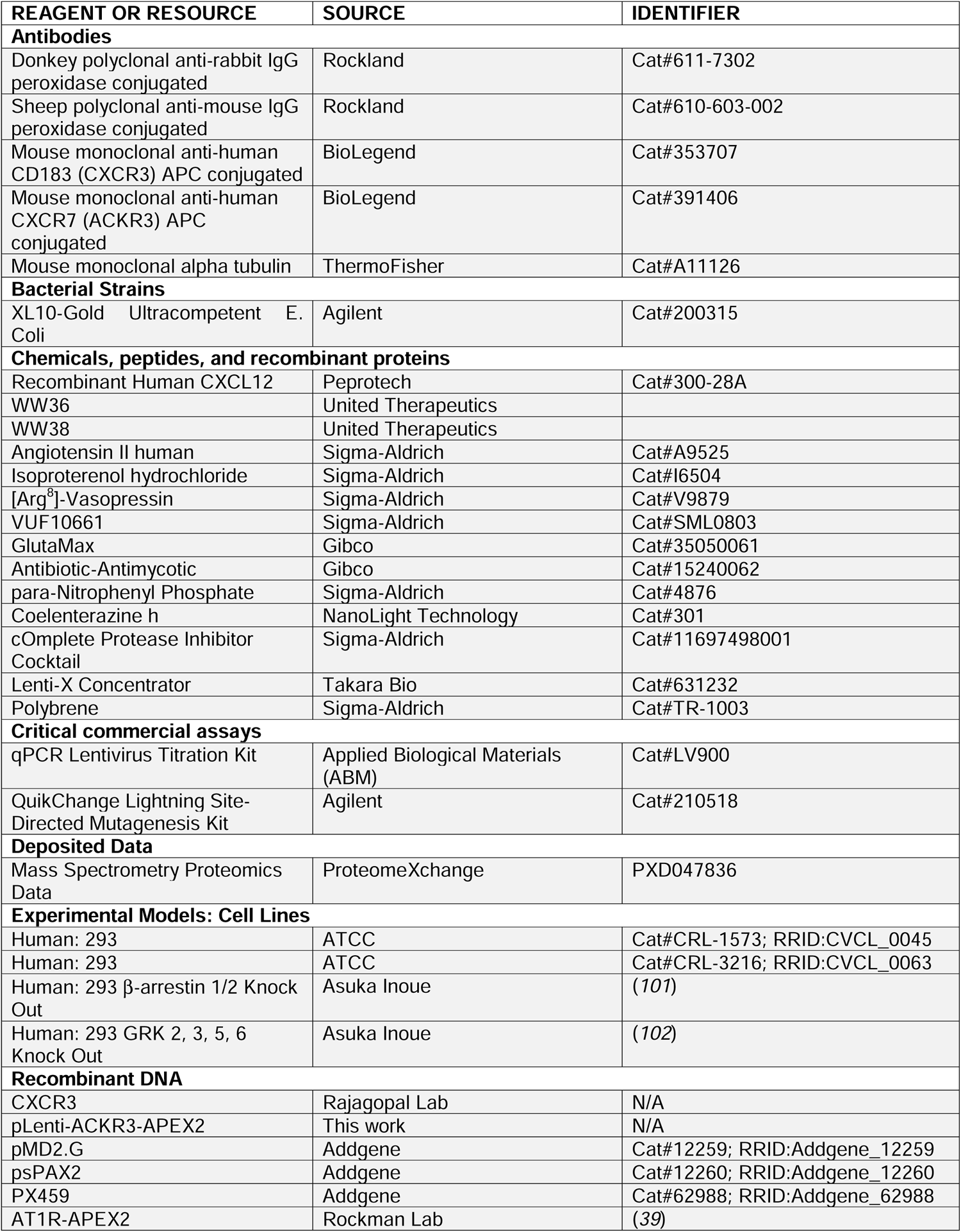
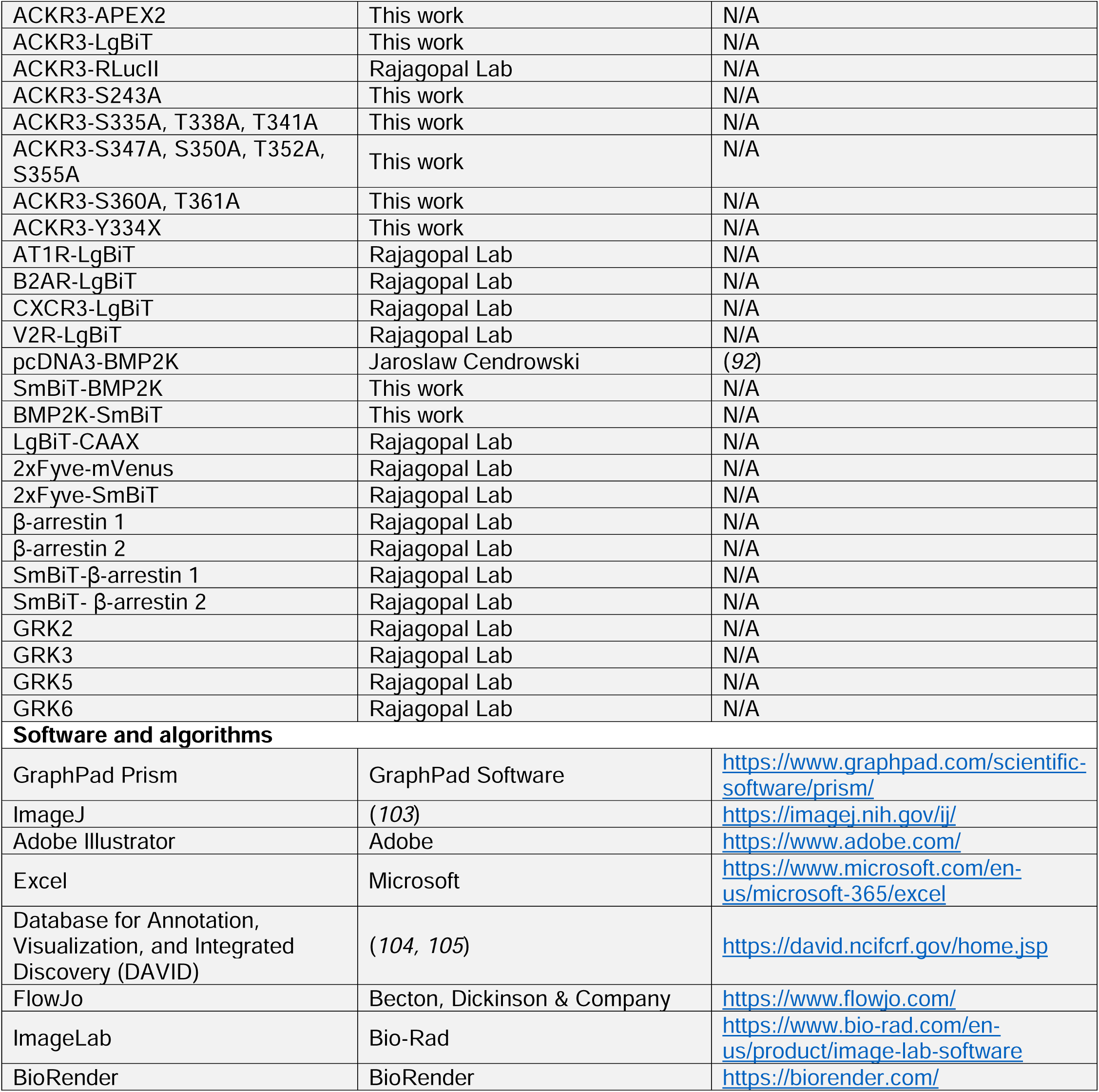

