## Supplementary figures and images for "ACKR3 Proximity Labeling Identifies Novel G protein- and β-arrestin-independent GPCR Interacting Proteins"

### Supplemental Figures

#### Supplemental Figure 1

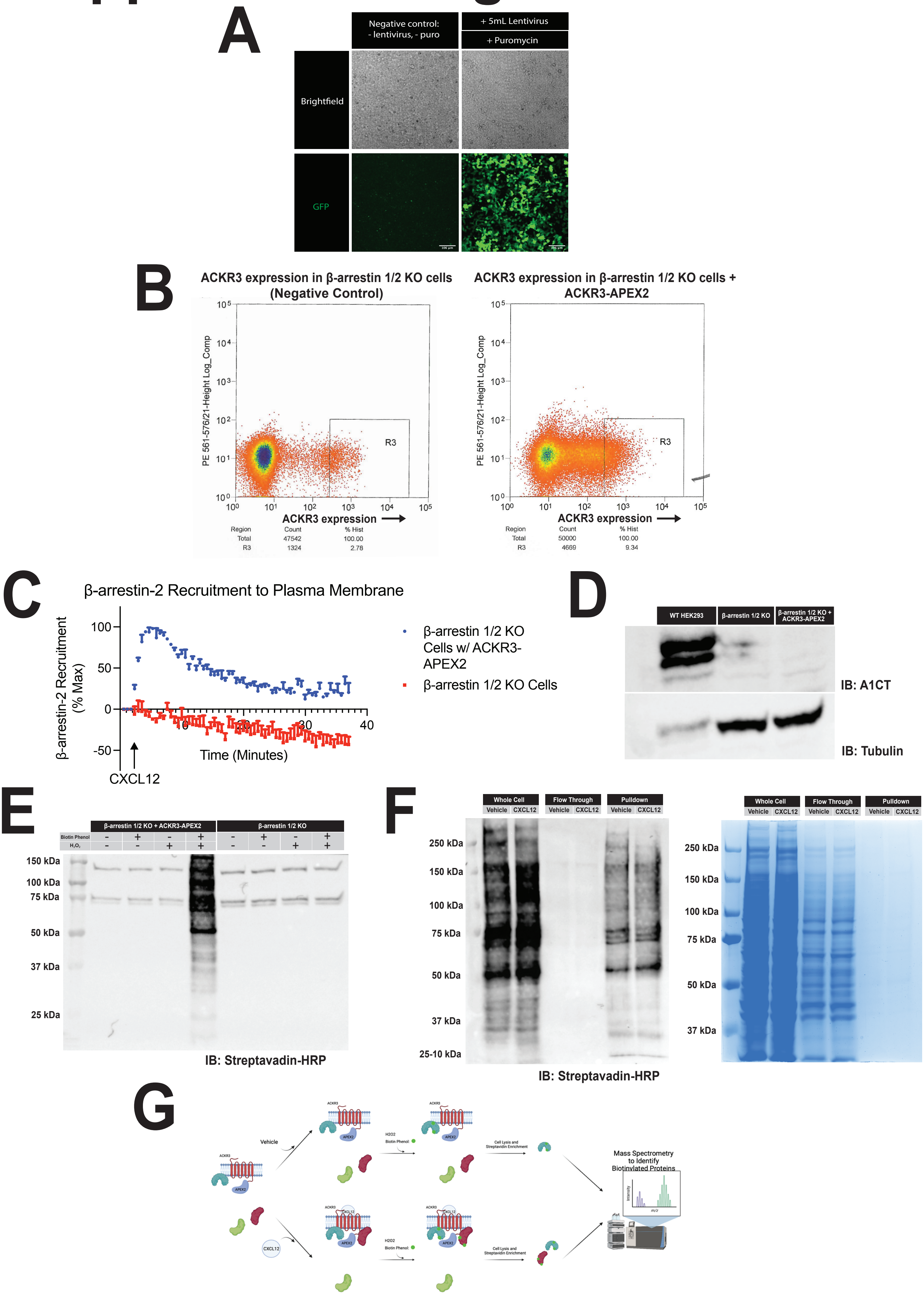

#### Supplemental Figure 2

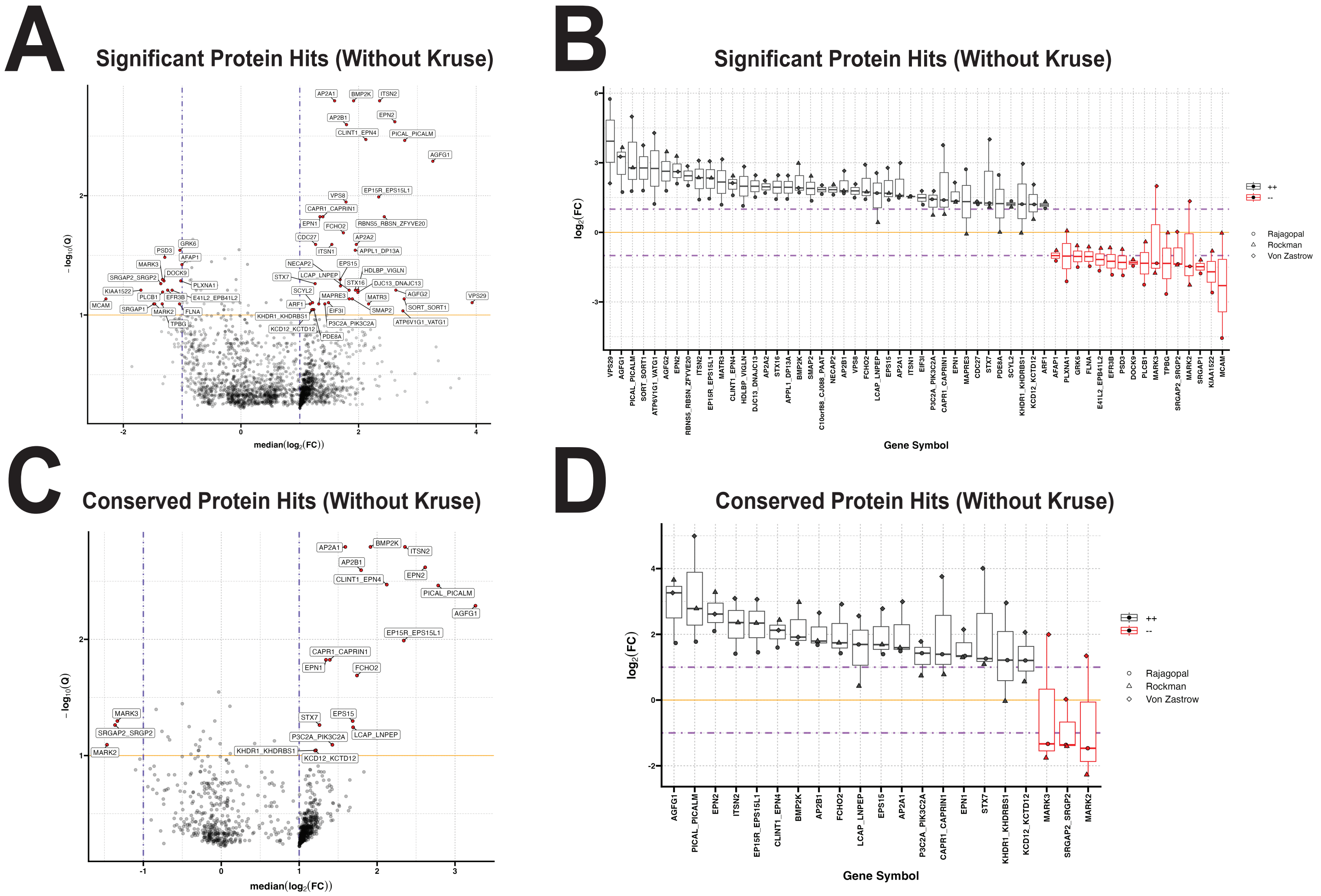

#### Supplemental Figure 3

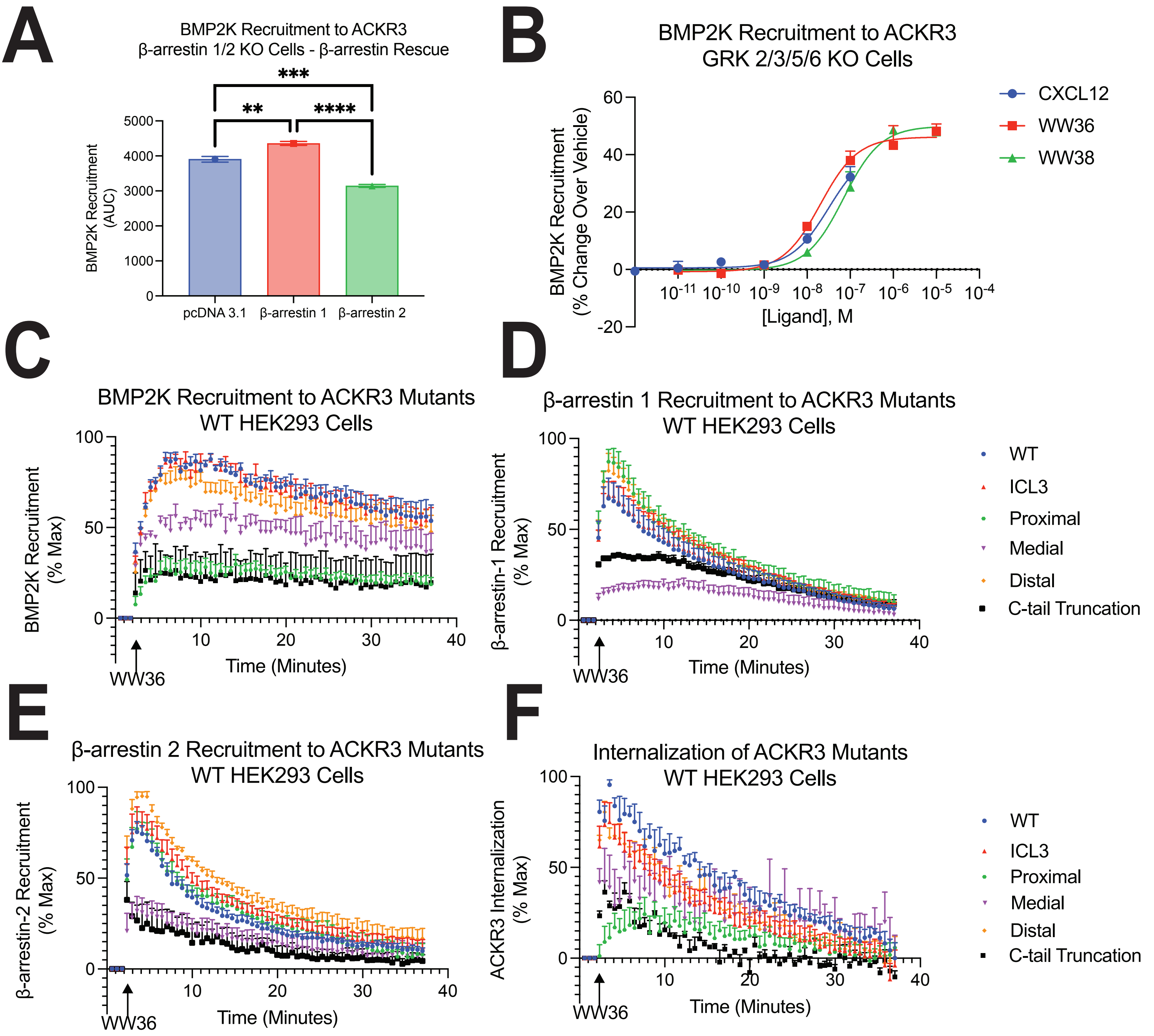
